# Genetic Mapping by Bulk Segregant Analysis in Drosophila: Experimental Design and Simulation-Based Inference

**DOI:** 10.1101/057984

**Authors:** John E Pool

**Affiliations:** Laboratory of Genetics, University of Wisconsin Madison, Madison, WI, 53706

**Keywords:** bulk segregant analysis, quantitative trait locus mapping, simulation, experimental design, *Drosophila*

## Abstract

Identifying the genomic regions that underlie complex phenotypic variation is a key challenge in modern biology. Many approaches to quantitative trait locus mapping in animal and plant species suffer from limited power and genomic resolution. Here, I investigate whether bulk segregant analysis (BSA), which has been successfully applied for yeast, may have utility in the genomic era for trait mapping in *Drosophila* (and other organisms that can be experimentally bred in similar numbers). I perform simulations to investigate the statistical signal of a quantitative trait locus (QTL) in a wide range of BSA and introgression mapping (IM) experiments. BSA consistently provides more accurate mapping signals than IM (in addition to allowing the mapping of multiple traits from the same experimental population). The performance of BSA and IM is maximized by having multiple independent crosses, more generations of interbreeding, larger numbers of breeding individuals, and greater genotyping effort, but is less affected by the proportion of individuals selected for phenotypic extreme pools. I also introduce a prototype analysis method for Simulation-based Inference for BSA Mapping (SIBSAM). This method identifies significant QTLs and estimates their genomic confidence intervals and relative effect sizes. Importantly, it also tests whether overlapping peaks should be considered as two distinct QTLs. This approach will facilitate improved trait mapping in *Drosophila* and other species for which hundreds or thousands of offspring (but not millions) can be studied.

## Introduction

Connecting phenotypic diversity to the genetic variants that encode it is a fundamental challenge for modern biology. In evolutionary research, there is strong interest in revealing the genetic architecture of adaptive phenotypic change, including the number of causative genes and mutations, and their functional and population genetic properties. In molecular genetics, the mapping of phenotypic differences from natural or induced mutations has great utility for elucidating genetic pathways that underlie specific biological processes. In animal and plant breeding, localizing the genes underlying agronomically important trait variation can be a key step toward genetic improvement.

Especially in species that can be experimentally crossed, quantitative trait locus (QTL) mapping provides an important tool for identifying genomic regions that contain causative genetic variants underlying a trait difference. Often, the F2 or later offspring of a cross between phenotypically contrasting parental strains are genotyped, individually or in groups, to identify sections of the genome that were inherited non-randomly with respect to the phenotype (often on the megabase scale). The simplest example of QTL analysis is F2 mapping, in which individual second generation offspring are phenotyped and genotyped. To achieve much genomic precision, however, this method requires the individual genotyping of a large number of F2 offspring. Preparing many genomic DNA libraries for next generation sequencing is often a time-and resource-intensive proposition, although progress has been made in this regard (Andolfatto *et al.* 2011).

Introgression mapping (IM) provides another alternative for QTL analysis. Here, following an initial cross between parental strains A and B, offspring of subsequent generations are repeatedly selected for strain A's phenotype, but are back-crossed to strain B (Figure 1). To allow recessive variants to be selected, this selection and introgression can be performed in every second generation. The desired result is an introgression line that is largely similar to strain B across the genome, but that matches strain A at loci that were selected along with the phenotype. A notable modern example of this approach is described by Early and Jones (2011), who introgressed a behavioral difference from *D. simulans* into *D. sechellia.* Here, 30 F2 females were tested for simulans-like behavior, and a subset was then back-crossed to *D. sechellia.* After repeating this process for 15 generations, next-generation sequencing was used to identify genomic regions that introgressed with the trait from *D. simulans.*

**Figure 1.**
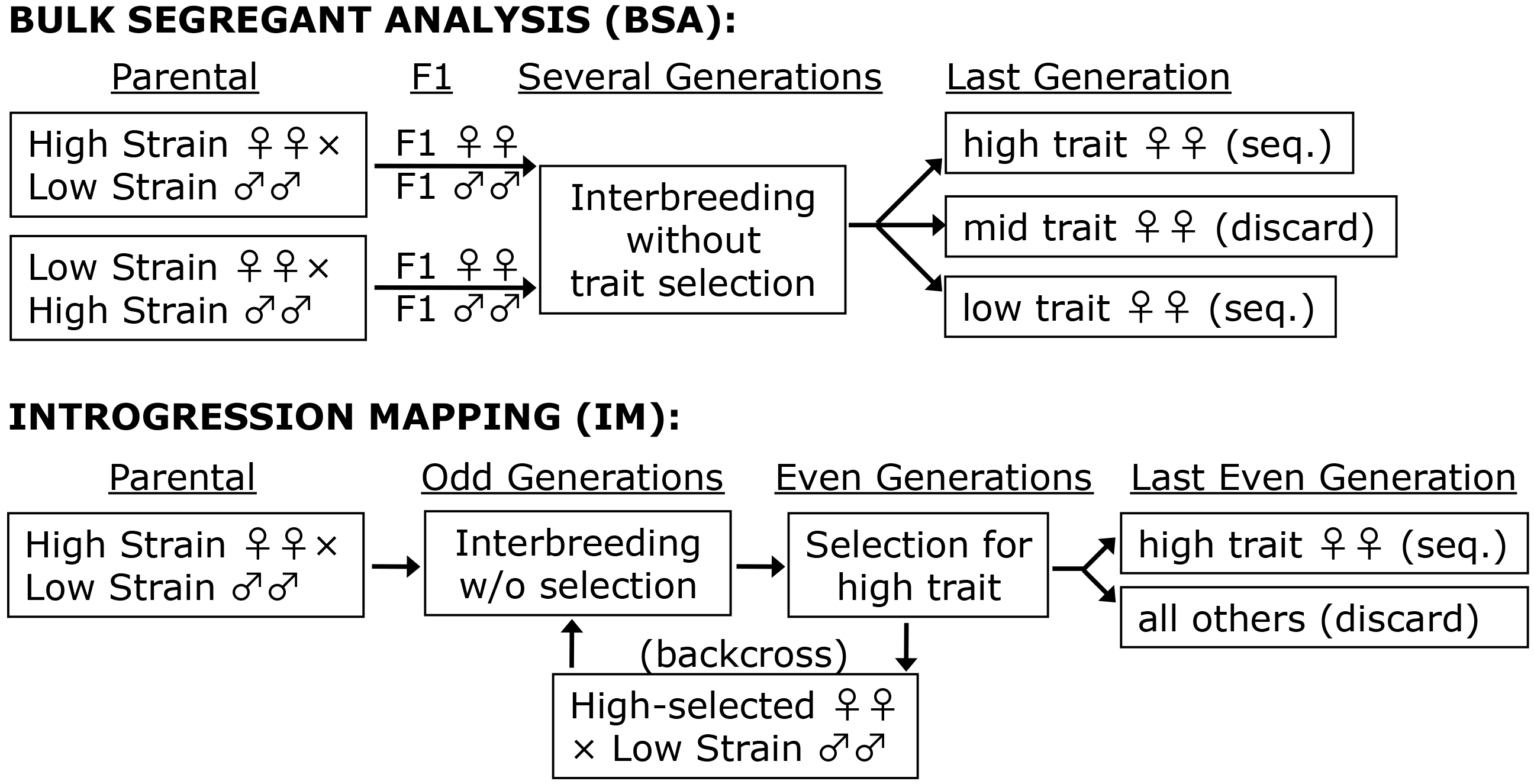
The investigated experimental designs for BSA and IM are illustrated. In BSA, offspring of reciprocal parental strain crosses are combined and allowed to breed without trait selection for a number of generations. Phenotyping occurs only in the final generation, and pools of individuals with the highest and lowest trait values are each sequenced. The IM framework investigated here involves trait selection and parental strain backcrossing every second generation (allowing recessive genotypes to be expressed). In the last generation, one phenotypic extreme is sequenced and compared against the backcross

In bulk segregant analysis (BSA), large numbers of progeny (from F2 or later generations) are sorted/selected by phenotype, then contrasting phenotypic pools of individuals are each genotyped (Figure 1) (Michelmore *et al.* 1991). Compared to IM, BSA may allow for a larger number of unique recombination events to be generated and sampled, which could yield sharper QTL peaks. Like IM, BSA does not require large numbers of offspring to be individually genotyped - instead each phenotypic extreme can be sequenced as a single pool. BSA has been applied very successfully for selectable traits in yeast *(e.g.* Ehrenreich *et al.* 2010; Magwene *et al.* 2011; Parts *et al.* 2011), facilitated by a small genome and the ease of generating millions of segregants. BSA has also seen diverse applications to trait mapping in multicellular organisms (e.g. Michelmore *et al.* 1991; Wicks *et al.* 2001; Baird *et al.* 2008; Van Leeuwen *et al.* 2012; Haase *et al.* 2015), including *Drosophila* (Lai *et al.* 2007).

Here, I use simulations to examine the mapping signals of BSA and IM under a wide range of experimental parameters for the mapping of multi-gene traits. I find that BSA produces stronger and better-localized mapping signals for all studied experimental designs. The tradeoffs of effort and performance indicated by these results, along with the new simulation programs that produced them, will help researchers design more effective mapping experiments.

I also use this BSA simulation approach to devise a new QTL inference method. Existing BSA analysis methods effectively identify QTLs from yeast data (*e.g.* Magwene *et al.* 2011; Edwards *et al.* 2012). However, these methods do not allow the discrimination of two nearby QTL peaks versus a single peak with noisy, ragged contours - an issue that may be more problematic for organisms in which many fewer segregants can be surveyed relative to yeast. These methods also do not estimate the relative strength of each QTL. The BSA inference method proposed here uses a multi-step simulation process to (1) identify significant QTLs and their genomic confidence intervals, (2) separate single from multiple linked QTLs, and (3) provide a rough estimate of the effect sizes of the identified QTLs.

This method is validated using simulations in the present study, and applied to data in an accompanying article (Bastide *et al.* 2016).

## Materials and Methods

### Preliminary Simulations for BSA and IM

Simulation programs were written to assess the QTL signals of BSA and IM (software related to this article is available at http://github.com/JohnEPool/SIBSAM1). BSA simulation analyses focused on a summary statistic, “ancestry difference” (*ad*). For a given genetic marker locus or genomic window of sequence, *a_d_* refers to the difference between the high and low phenotypic pools in the proportion of ancestry from the parental strain with the higher phenotypic value. For example, if the high phenotypic pool is estimated to have 60% of its ancestry from this parental strain at a particular locus, and the low phenotypic pool 40%, then *a_d_* = 0.6 − 0.4 = 0.2. For IM, the proportion of ancestry in the mapping population from the non-backcross parental strain (*a_p_*) was evaluated. This quantity may approach zero for non-causative loci after many generations of back-crossing to the other parental strain. For each statistic, I examined how often the tallest local QTL peak was observed within 0.5 centiMorgans (cM) of the true simulated target locus, and the average (median) distance between the QTL peak and the target locus.

The BSA and IM simulators are largely similar. These programs track parental strain ancestry along the chromosomes of each individual in the mapping population, from the F1 generation until the end of the experiment. A Poisson-distributed number of recombination events happen each generation, with the expected number for each chromosome being its length in Morgans (interference is not modeled). To focus on the case of *Drosophila,* chromosomes X, 2, and 3 were explicitly simulated, and no recombination was allowed in males. A total of 5,000 markers/windows were simulated on each chromosome. In the BSA simulation, a specified number of individuals exist in each new generation, and each one draws random parents from the previous generation, with no phenotypic selection until the last generation. In the IM simulations, individuals were subject to phenotypic selection in every second generation.

Phenotypes for each individual were modeled based on genotypes and random variance (the latter may stem from environmental effects, measurement error, or other causes). For most of these preliminary simulations, the same number of equal-effect loci were simulated on each chromosome arm (X, 2L, 2R, 3L, 3R). Random variance was added by modifying each individual' s phenotypic value by a normally-distributed random effect with mean zero and standard deviation (SD) equal to the average trait value. For example, if each of the five arms holds a single QTL that adds 1 to a diploid individual' s phenotypic value for each allele inherited from the high parental strain, the range of genetic contributions could range from 0 to 10, with a mean of 5, and the SD for environmental variance would also be 5. Phenotypic selection was then based on choosing a defined quantile (*q*) of individuals from the mapping population with the highest and the lowest phenotypic values.

For BSA, phenotypic selection happens only at the end of the experiment, followed by sequencing/genotyping of both high and low phenotypic pools. For IM, the last batch of selected individuals is sequenced and compared against the parental strains. The simulations model “depth” of sequencing coverage (or genotype sampling), drawing an appropriate number of random ancestry-informative reads from the selected pool of individuals for each window/marker. The proportion of ancestry from each parental strain is then calculated, and thus depends on both the sampling of individuals and the sampling of sequence reads.

To facilitate consistent analysis, QTLs in these preliminary simulations were spaced uniformly and each was assigned a specific analysis zone along the chromosome. For example, if the X chromosome had five QTLs, they would be placed at relative positions 0.1, 0.3, 0.5, 0.7, and 0.9 (representing the chromosome as a 0 to 1 interval). Their zones of analysis would then be 0 to 0.2, 0.2 to 0.4, and so on. The assessment of QTL signal strength and precision was based on the location within its zone of the highest QTL peak *(i.e.* the maximum *a_d_* or *_d_*), relative to the true QTL position.

Most simulation analyses assumed that each mapping experiment would be analyzed separately, However, I also investigated cases where multiple independent mapping populations were constructed from parental strains sharing the same causative genetic differences. Here, *a_d_* or *a_p_* for each window was summed across replicated mapping populations.

For a wide variety of experimental parameter combinations, 1,000 independent replicates were simulated and analyzed, and statistical performance was compared between these scenarios to aid in the optimization of experimental design.

### Simulation-Based Inference of QTL from BSA: Overview

Preliminary empirical BSA data from the Pool laboratory indicated the need for a QTL inference method capable of dealing with neighboring QTLs that have wide, overlapping statistical signals. Such scenarios are difficult to account for in most analysis approaches, but the simulation framework described above offers a potentially flexible foundation for QTL inference. I therefore developed a method of “Simulation-based Inference for Bulk Segregant Analysis Mapping” (SIBSAM). SIBSAM uses BSA simulations analogous to those described above, in order to identify and localize significant QTLs, estimate their strength, and distinguish individual QTL among clusters of linked causative loci.

Throughout the SIBSAM pipeline, the distinction between primary QTL peaks and secondary QTL peaks is relevant. A primary QTL peak is defined based on the highest value
of *a_d_* across a continuous interval in which this statistic remains above zero (which is the null value expected in the absence of causative loci). A secondary QTL peak within that same interval has a lower height than its associated primary peak. An important quantity in assessing the significance of a secondary peak is its “secondary deviation” (*v*), defined as the difference between secondary peak height and the minimum *a_d_* value between the primary and secondary peaks (Figure 2). Multiple secondary peaks may be associated with the same primary peak, impacting the calculation of *v*, as discussed below.

**Figure 2.**
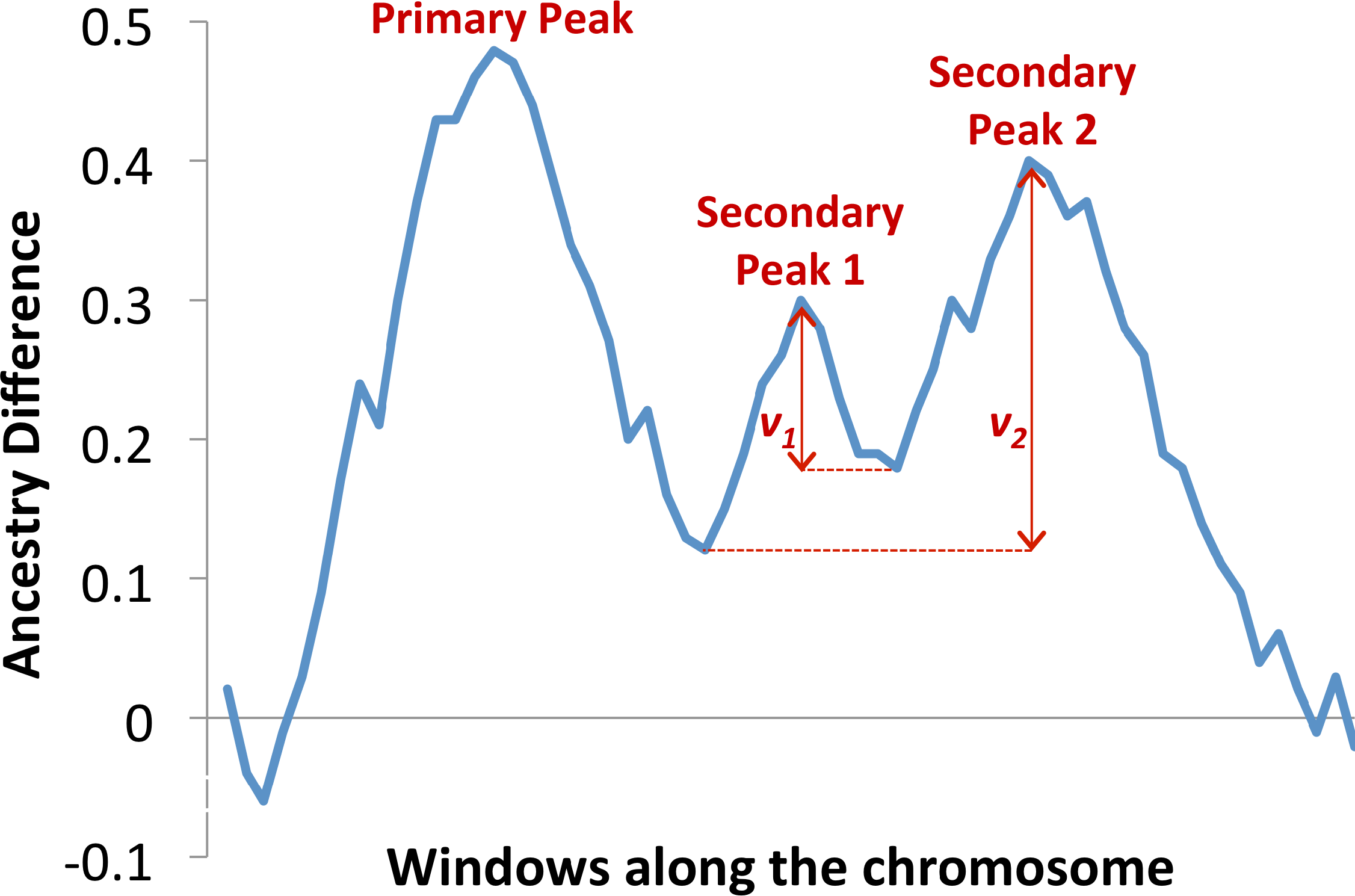
Definitions of primary and secondary peaks, along with secondary deviation, used by SIBSAM are illustrated here. Among a group of contiguous windows with smoothed *a_d_* values above zero, the primary peak is defined by the window with the highest value. Secondary peaks represent other local maxima, and their significance is judged based on secondary deviation (*v*). Secondary deviation is determined by the difference in *a_d_* between the secondary peak's maximum value and the minimum value between that peak and the primary peak (or a taller secondary peak, whichever minimum is greater).

A schematic of the SIBSAM pipeline is illustrated in Figure 3. First, primary and secondary peaks of *a_d_* are identified from the empirical data. To determine which primary peaks are unexpected in the absence of true QTLs, null simulations are conducted in which phenotypes are determined by non-genetic factors only. *P* values can then be obtained for each primary peak. Next, simulations with a single causative QTL are conducted. Based on a rejection sampling approach, estimates of the strength and genomic confidence intervals of each significant primary peak are obtained, along with a *P* value for each secondary peak. Lastly, simulations involving a cluster of linked QTLs are conducted, reflecting a primary peak and its associated secondary peak(s). This phase allows for the refinement of strength estimates and genomic confidence intervals for each peak in the cluster.

**Figure 3.**
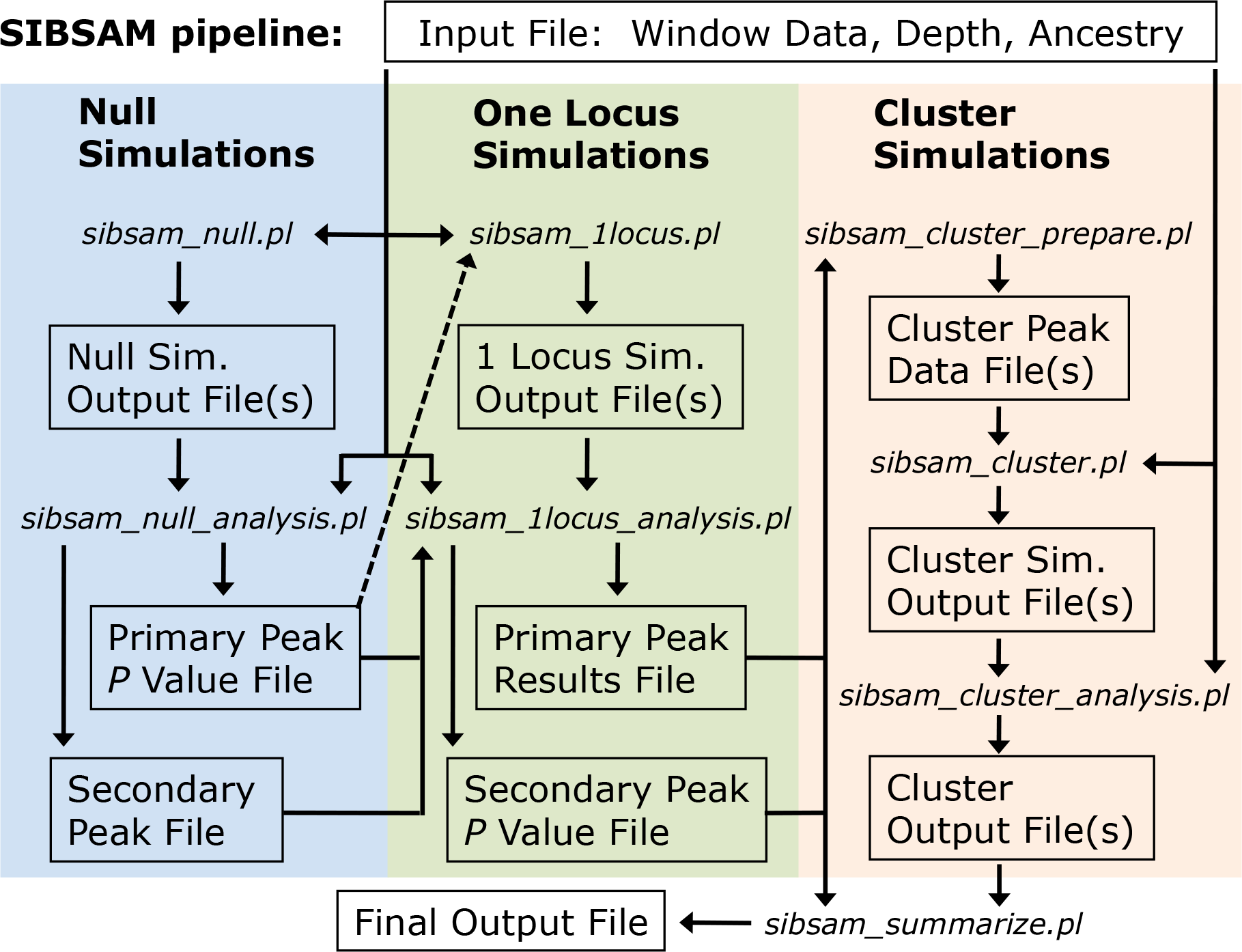
A flow chart illustrating the SIBSAM analysis pipeline is shown. A single input file contains physical and genetic map positions of window boundaries for all chromosomes, along with ancestry difference values and informative depth (the number of reads within information about parental strain ancestry) for each window. Null simulations with no true QTLs are used to identify significant primary peaks in the empirical data. Simulations with one QTL (matching a primary peak location) are then used to estimate confidence intervals for primary peak effect size and genomic location, while also identifying significant secondary peaks. For any primary peak with significant secondary peaks, cluster simulations are conducted with QTLs at each peak's location, in order to generate final confidence intervals for effect size and genomic location. These analyses are summarized into a single output file containing all relevant inferences for each significant peak.

All of the above simulations operate with user-defined windows of variable cM length. These windows could also be viewed as markers separated by various cM distances, but this article's terminology mainly assumes that QTL mapping data comes from the full resequencing of mapping population genomes. In the examples presented here, the window bp spans were based on *D. melanogaster* polymorphism data (Lack *et al.* 2015) and cM distances were calculated from empirical recombination rate estimates (Comeron *et al.*2012). Windows were defined to each contain 200 non-singleton variable sites from the Zambia-Siavonga population sample. The user can also define the “informative depth” for each window in each phenotypic pool. This quantity refers to the number of sequence reads that contain information about parental strain ancestry. The simulator will draw a corresponding number of alleles at this window for ancestry proportion calculations.

### SIBSAM Identification of Primary and Secondary Peaks from Empirical Data

Primary and secondary peaks of *a_d_* are identified from data based on preliminary thresholds for primary peak height and secondary peak deviation *(a_dt_* and *V_t_*, respectively), plus an optional smoothing step. The two thresholds should represent values low enough that no shorter peak would be statistically significant (the default value for both is 0.1). The smoothing enabled here is a simple weighted average. On each side of the focal window, *m* flanking windows are included (the default used here is *m* = 4). The focal window receives a weight of *m* + 1, the adjacent window on each side receives a weight of m, the next windows receive a weight of *m* − 1, and so on until the *m*th window to each side receives a weight of 1. Alternative smoothing schemes are not a focus of this study; the optimal strategy should depend on the data being analyzed. Empirical and simulated *a_d_* values must be smoothed using the same procedure.

Primary peak identification is straightforward: the highest value of *a_d_* in a continuous block of windows with *a_d_* > 0, conditional on the peak value of *a_d_* exceeding *a_dt_.* To identify secondary peaks, local minima and maxima of *a_d_* moving away from the primary peak are noted. A recovery, beyond *V_t_*, from the low point since the last peak signifies a new secondary peak. When *a_d_* drops more than *V_t_* below this secondary peak's maximum value, this peak ends and its maximum value and associated window position are noted. Statistical significance of these primary and secondary peaks, along with their confidence intervals and relative strengths, will be assessed in subsequent stages of this pipeline.

### SIBSAM Identification of Significant Primary Peaks

The false positive probability (*P*) for each primary peak is estimated by comparing empirical peak heights against simulations under the null hypothesis of no true QTLs, in which all phenotypic variance in the mapping population is random with respect to genotype. All primary peaks exceeding *a_dt_* from each simulation replicate are noted. The enrichment (*e*) of peaks equal to or greater than a given peak's height in the real data is given by the ratio of the frequency of peaks of this height in the real data relative to the simulated data. If there is an enrichment (*e* > 1), an estimate of the proportion of real peaks of this height representing false positives is then given by 1 / *e*. For example, if *a_d_* peaks of at least 0.2 in height are three times more common in the empirical data than in null simulations, then on average one out of three such empirical peaks can be explained by the expected false positive rate. Primary peaks with an estimated *P* less than some threshold (by default, 0.05) are carried forward for subsequent analysis.

### SIBSAM Inferences from Single QTL Simulations

Genomic simulations with a single QTL are used to estimate the genomic confidence intervals and strength of each primary peak, along with a *P* value for each secondary peak. Single QTL simulations are performed with each fixed genomic positions corresponding to the window with the peak maximum *a_d_* for each QTL, thus conserving local window patterns of depth and cM distance. For a given set of simulated genomes from the mapping population (pre-selection), a random QTL effect size is drawn. Such a QTL is then separately simulated at each position corresponding to an empirical primary peak, with phenotype simulation and read sampling performed separately in each case. The simulated cage ancestries are reused for each separate QTL simulation as a time-saving efficiency.

The simulated QTL strength, s, ranging from 0 to 1, is the estimated proportion of variance that a QTL explains among the mapping population individuals. In these single locus simulations, all other phenotypic contributions are modeled as random variance, which here is intended to encompass the effects of unlinked QTLs in addition to non-genetic effects on phenotypic measurements. The amount of random variance simulated is fixed to approximate the variance contributed by a codominant locus in which each allele adds 1 to the phenotypic score. This effect was implemented by obtaining Gaussian random values with mean 0 and standard deviation 1, and then multiplying each value by 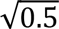 to obtain the random variance effect on each individual's phenotypic score. The simulated effect size of each QTL, e, describes the quantity that each allele of this locus (inherited from the high parental strain) adds to an individual's phenotypic score. Since random effects correspond to the variance contributed by a locus with *e* = 1, the proportion of variance contributed by a single QTL (*s*) is equal to *e* / (1 + *e*). And correspondingly, a single QTL intended to have strength *s* is simulated with an effect size *e* = *s* / (1 + *e*).

For each simulated replicate, the simulated strength is recorded, along with each QTL's maximum height, peak window location, and maximum secondary deviation. To analyze the one locus simulation data for each primary peak, a rejection sampling approach is used to identify simulation replicates in which maximum *a_d_* falls within a specified tolerance (default 0.025) of the empirical peak's maximum *a_d_.* For each accepted simulation replicate, the strength of the simulated locus goes into the posterior distribution for the empirical QTL's strength (from which strength values corresponding to the 0.05, 0.5, and 0.95 quantiles are returned). A genomic confidence interval is similarly obtained by examining the far left and far right quantiles for the simulated peak locations resulting from a QTL simulated at the empirical peak location. This assumes a certain transitivity. Here, we are simulating QTLs with fixed positions and observing how far away the maximum *a_d_* falls in these simulations. In the empirical data, we observe the location of the maximum *a_d_*, and we'd like to know how far from this window the true QTL might be. Thus, we assume the distances from true QTL to maximum *a_d_* in the simulated data are a good proxy for the distances between maximum *a_d_* and true QTL in the empirical data.

Lastly, the secondary deviations from each accepted simulation enable *P* values to be calculated for each of the empirical primary peak's associated secondary peaks. If more than one secondary peak is present on the same side of the primary peak in the empirical data, the tallest secondary peak is tested first, and its *v* is based on the difference between its height and the lowest *a_d_* value between itself and the primary peak (even if other secondary peaks exist between this peak and valley; Figure 2). For a shorter secondary peak between a primary peak and a taller secondary peak, *v* would be defined as the difference between its height and the higher of the valleys on either side of it. Giving taller peaks this priority avoids the situation of a shorter secondary peak being deemed significant and a taller peak beyond it missing this threshold (as might occur if secondary peaks were simply evaluated sequentially by position). After such adjustments, each secondary peak deviation in the empirical data associated with this primary peak is compared to the distribution of *v* from accepted simulations. The proportion of simulations with a *v* greater than observed for a given empirical secondary peak becomes the *P* value for that peak (*i.e*. the probability of getting a secondary deviation this extreme when the true model is a single QTL of the observed magnitude).

### SIBSAM Inferences from QTL Cluster Simulations

In cases where an empirical primary peak is accompanied by one or more statistically significant secondary peaks, the strengths and confidence intervals of all peaks in this “QTL cluster” are best approximated from simulations that include each member QTL. For example, a pair of nearby QTLs may each add to the *a_d_* peak height of the other, leading to overestimates of effect size. Therefore, multi-QTL simulations are conducted separately for each QTL cluster inferred from the empirical data. For simplicity, the window position of each simulated QTL is fixed according to the windows showing maximum *a_d_* for each significant peak in the empirical cluster. To examine each QTL separately, each is assigned an analysis zone with boundaries corresponding to the empirical valleys (local minima) between peaks. Moving away from the outer peaks in the cluster, this analysis zone is bounded only by the ends of the chromosome.

For each cluster simulation replicate, a random strength value is first drawn for the full cluster (representing the cumulative proportion of phenotypic variance explained by the QTLs in this cluster). That cluster strength is randomly apportioned among the QTLs, and each peak's strength is then translated into the simulated effect size as described above.

A cluster simulation replicate is accepted only if the local maximum *a_d_* in every QTL's analysis zone falls within a tolerance of the corresponding empirical peak heights. Here, it could be necessary to use a slightly higher tolerance value to accrue enough accepted simulations (default *a_d_* tolerance 0.05). This or any other simulation step in SIBSAM can be parallelized to increase the number of replicates, followed by joint analysis of multiple simulation output files (Figure 3).

The estimated strength of each peak in cluster, along with confidence intervals of strength and genomic position, are obtained from a similar rejection process as described for the one locus simulations (based on the distribution of strength values and peak locations for that peak among the accepted simulations). Thus, the cluster QTL simulations provide estimates of effect size and genomic confidence intervals for all significant secondary peaks. They also replace prior estimates of these quantities for the associated primary peaks, since cluster estimates that account for the effects of linked QTLs should be more accurate.

The final SIBSAM output file contains, for each significant primary and secondary peak, its *P* value, the genomic coordinates of the peak window and the confidence interval for the QTL's genomic location, and the point estimate and confidence interval for QTL strength. Information such as *P* values for non-significant peaks can be found in the intermediate files produced at different stages of the SIBSAM pipeline (Figure 3).

### Simulations testing the performance of SIBSAM

Simulation testing of SIBSAM was performed to test its QTL detection power under different scenarios, and to confirm that estimates and confidence intervals of genomic location and QTL strength were performing in line with expectations. Although a nearly infinite range of scenarios could potentially be investigated, I focused on experimental parameters relevant to our current empirical applications in *Drosophila* (*e.g*. Bastide *et al.* 2016), in which 1,200 individuals interbreed for 16 generations, and 10% phenotypic tails are selected for sequencing. Test simulations sampled 1,000 informative reads for each window for each phenotypic pool, which is about half the median depth per window from current empirical applications (*e.g*. Bastide *et al.* 2016). Windows were designed to each contain 200 non-singleton variable sites in the Zambia-Siavonga population genomic data described by Lack *et al.* (2015). These 14,107 windows had a median length of 6.8 kb.

Simulations with one genuine QTL were performed with varying locus strengths (*s* = 0.05, 0.1, 0.15, 0.2, 0.25, 0.33, 0.5). These initial test simulations used fixed genomic positions corresponding to the locations of *Drosophila* pigmentation genes *tan* (on the X chromosome) and *ebony* (on arm 3R). Additional 3R scenarios with *s* = 0.2 investigated the consequences of the remaining variance being due to unlinked QTLs (1 with *s* = 0.8 or else 4 others with *s* = 0.2) instead of random Gaussian variance. Comparing each test replicate against SIBSAM null simulations revealed the true positive rate for QTL detection. Running the test replicates through the SIBSAM one locus simulation analysis indicated the frequency at which secondary QTLs were falsely inferred, along with allowed the inferred distributions of QTL strength and genomic location to be compared against known true values.

Additional simulations were conducted (focusing on the 3R location) to investigate SIBSAM's performance in the presence of two linked QTLs. Scenarios with symmetric QTL strength (s = 0.15 or 0.3) and asymmetric QTL strength (s = .15 and 0.3) were investigated. The distance between the two QTLs was varied at 2.5, 5, 10, and 25 cM. The test replicates were then evaluated with SIBSAM to (1) test the power to detect one or both QTLs, (2) test the rate of falsely detecting three or more QTLs, (3) evaluate the performance of QTL localization, and (4) evaluate the performance of QTL size estimation.

## Results

### Initial Simulation Study of BSA and IM

Simulations were performed to examine the properties of QTL signals under BSA and IM approaches. Importantly, these exploratory simulations are not connected to any formal QTL inference. Instead, they focus on the performance of summary statistics related to the signature of a QTL. For BSA, I examine ancestry difference (*a_d_*), the difference between high and low phenotypic pools in the proportion of ancestry sampled from the parental strain with the higher phenotypic value (at a particular genomic locus). For IM, I examine ancestry proportion (*a_p_*), the proportion of the mapping population's ancestry that derives from the non-backcross parental strain. Rather than focusing on the raw values of these statistics, I assess the performance of BSA and IM by examining the genetic distance between a true simulated QTL and the “QTL peak” (the maximum value of *a_d_* or *a_p_* in this part of the genome).

The above approach allows a wider range of scenarios to be examined than would be computationally feasible under the full SIBSAM inference process. Beyond a tentative comparison of the genomic precision of BSA vs. IM, an important goal here is to optimize critical experimental parameters to improve the outcomes of future trait mapping studies.

As a point of reference, these simulations began with a “default” scenario in which 600 individuals were bred each generation, for 10 total generations, phenotypic selection retained the 20% most extreme individuals in each direction, and each window/locus had a sequencing depth of 300. Individual parameters were then varied, alone or in combination, and the accuracy of the *a_d_* or *a_p_* signal was examined.

First, performance was examined when tandemly varying the number of QTLs and the number of independent crosses. Within each simulation case, all QTLs were of equal magnitude and explained 5/6 of total phenotypic variance. Independent crosses were simulated under the assumption that all pairs of parental strains share a given QTL difference between them. When multiple crosses were analyzed together, *a_d_* or *a_p_* were added between crosses for each genomic window to test whether a more precise localization emerged from this joint signal. Three primary themes emerged from this analysis. First, BSA outperformed IM for any given combination of crosses and loci (Figure 4). Second, combining data from multiple crosses had a markedly positive effect on the accuracy of these ancestry signals. Third, performance showed a predictable decline for more/weaker QTLs. Still, cases with multiple crosses still managed relatively stronger performance for more polygenic scenarios (Figure 4), particularly in the case of BSA. For simplicity, the remaining simulations below will focus on a single cross replicate and a scenario with five QTLs.

**Figure 4.**
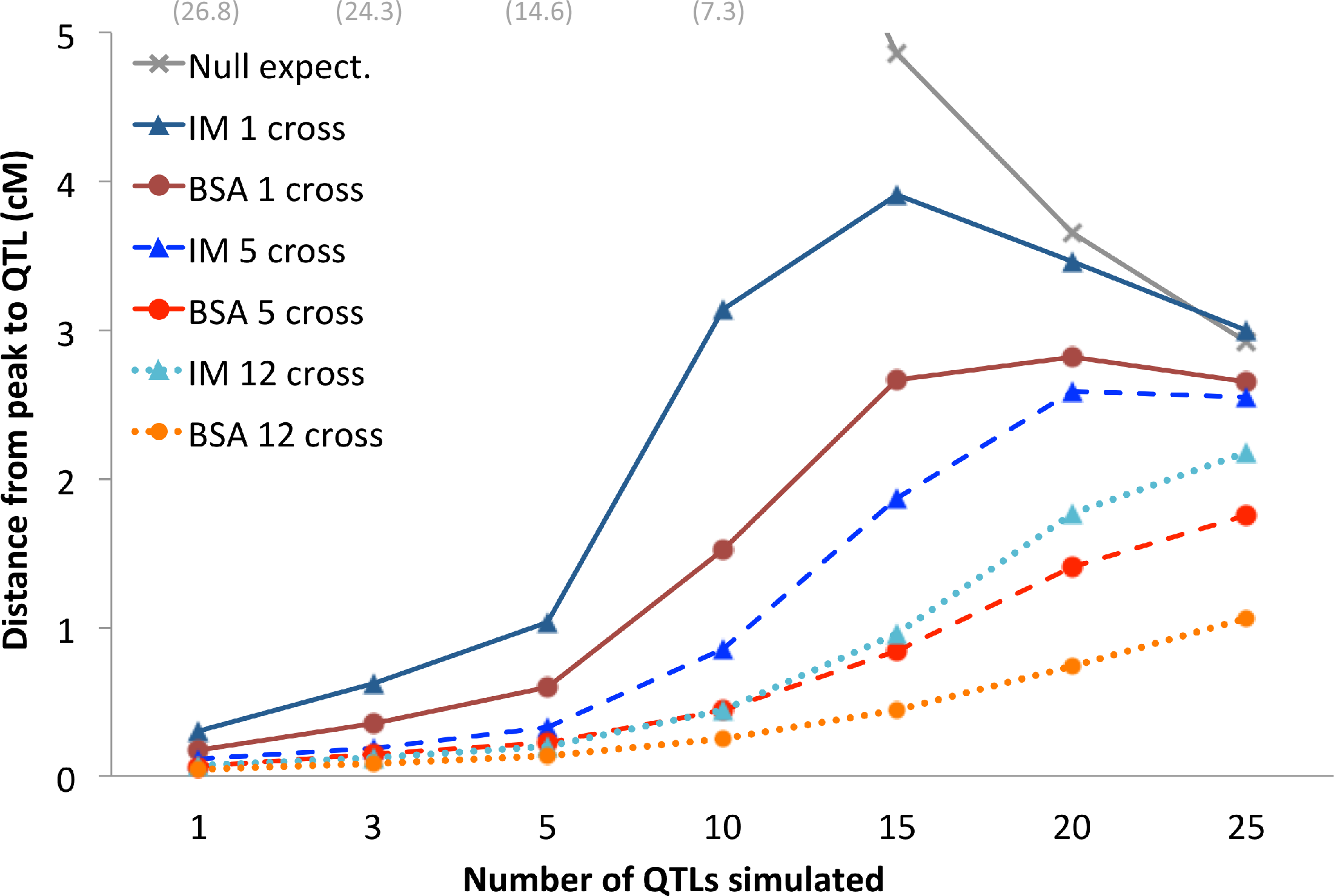
Results are shown for exploratory BSA and IM simulations with varying numbers of QTLs and numbers of jointly-analyzed independent crosses. As a proxy for method performance, the median centiMorgan distance between the true QTL and the statistic maximum (of *a_d_* for BSA or *a_p_* for IM) is shown. The null expectation for a randomly located peak within a QTL's analysis window is also shown (gray). These results indicate: (1) the increasing challenge of more polygenic scenarios for all approaches, (2) a general advantage of BSA over IM, and (3) the utility of combining data from independent crosses that all share a given QTL in common.

The number of generations before genotyping/sequencing was also varied. Strong performance improvement was observed by increasing the number of generations to 8 or 10, with further increases yielding ongoing but diminishing improvements (Figure 5A). Additional generations allow more recombination between parental genetic backgrounds, which should lead to sharper QTL peaks.

**Figure 5.**
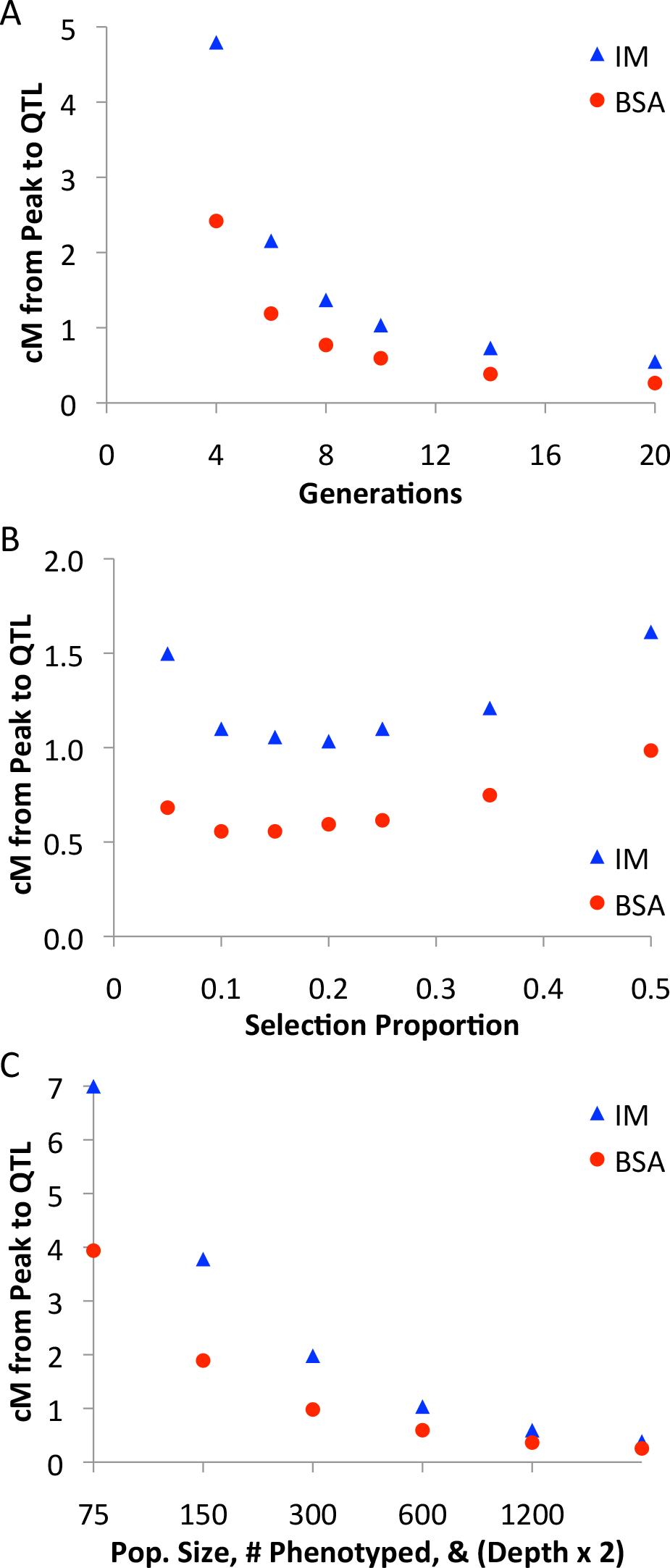
The results of exploratory BSA and IM simulations are shown in which one c more experimental variables were manipulated. (A) Increasing the total number of generations in the experiment reduces the median centiMorgan distance between the 1 QTLs and the observed peak. (B) A broad optimal range of selection proportion exists the focal BSA and IM scenarios. (C) Scaling up the experimental population size (and h the number of phenotyped individuals), along with the sequencing depth, leads to improved statistical performance.

Past results indicate that selecting only the most extreme individuals is not optimal for BSA (Magwene *et al.* 2011). Concordantly, for the focal simulation scenario studied here, optimum bulk proportions were around 10-15% for each BSA pool, and 20% for the single IM pool (Figure 5B). These results appear to reflect a balance between enriching for causative genotypes (favoring fewer individuals) and minimizing the effects of random sampling variance (favoring more individuals). Thus, both BSA and IM studies may benefit from selecting significant numbers of individuals, which should help to maximize the diversity of recombination breakpoints represented in the final data.

Related to the issue of sampling variance are parameters such as the number of individuals present in each generation and the number of genotypes sampled in the data (*e.g*. sequencing depth). When simulations jointly scaled up the number of individuals present in each generation, the number sampled for sequencing, and the sequencing depth, performance improved considerably (Figure 5C). The number of individuals sampled in the final generation made a particular difference, at least if depth was scaled up linearly (Figure S1). Increasing sequence depth consistently led to better performance (via a reduction in sampling variance), although with some diminishing returns (Figure 6).

**Figure 6.**
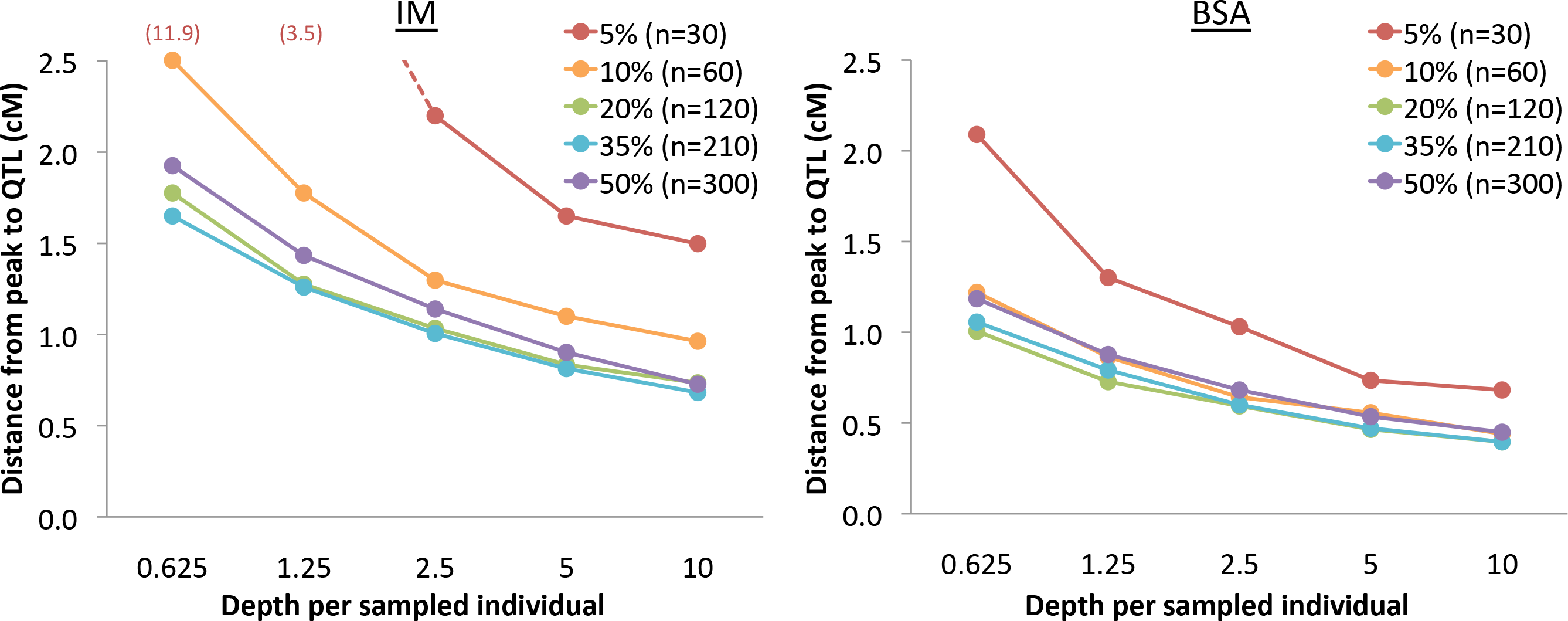
Outcomes of exploratory BSA and IM simulations with variable sequencing depth are shown. To more clearly illustrate the influence of depth on sampling variance, depth is plotted in terms of the average number of reads for each individual in a phenotypically selected pool. From a group of 600 phenotyped individuals, results for a series of selection proportions are illustrated. Results illustrate the advantage of increased sequencing depth, with some diminishing returns.

Simulations also considered the interaction between selection proportion and population size. The optimal selection proportion (*s*) tends to scale inversely with population size (*N*). For BSA population sizes between 100 and 2,400, there was a relative stability in the optimal number of sampled individuals for sequencing (*N_s_*), with this quantity ranging only from 35 to 60 (Table S1). In line with the findings of Magwene *et al.* (2011), this result suggests that reducing sampling variance is of primary importance, whereas enriching for the most phenotypically extreme individuals is a secondary priority.

### Simulation Testing of the SIBSAM Pipeline

As elaborated in the Materials and Methods section, I developed a prototype method for Simulation-based Inference for BSA Mapping (SIBSAM). The flexibility of this simulation-driven pipeline allows a range of inferences, including for challenging cases in which two or more QTLs are part of the same complex peak (Figure 2). The goals of SIBSAM include assessing the significance of peaks, and estimating the strength and genomic confidence interval of significant QTL. The performance of SIBSAM was assessed via a series of test simulations with one or more QTLs. While a vast range of QTL and experimental scenarios could potentially be examined, I focus here on parameters relevant to ongoing empirical work in *Drosophila* (Bastide *et al.* 2016). The BSA experimental design simulated here went for 16 generations, with 1,200 individuals in each generation, with 600 females phenotyped in the last generation with 10% pools selected, and 1,000 informative sequence reads for each genomic window.

For the above scenario, SIBSAM's QTL detection power went from weak for a QTL explaining 10% of the experimental population's phenotypic variance (with the remainder due to random environmental or measurement variance) to strong for a 20% QTL, with intermediate power for 15% QTL (Figure 7). As illustrated by the exploratory simulations above, the performance of QTL mapping is likely to be improved by increasing the number of generations, the population size, sequencing depth, and/or the number of independent crosses.

**Figure 7.**
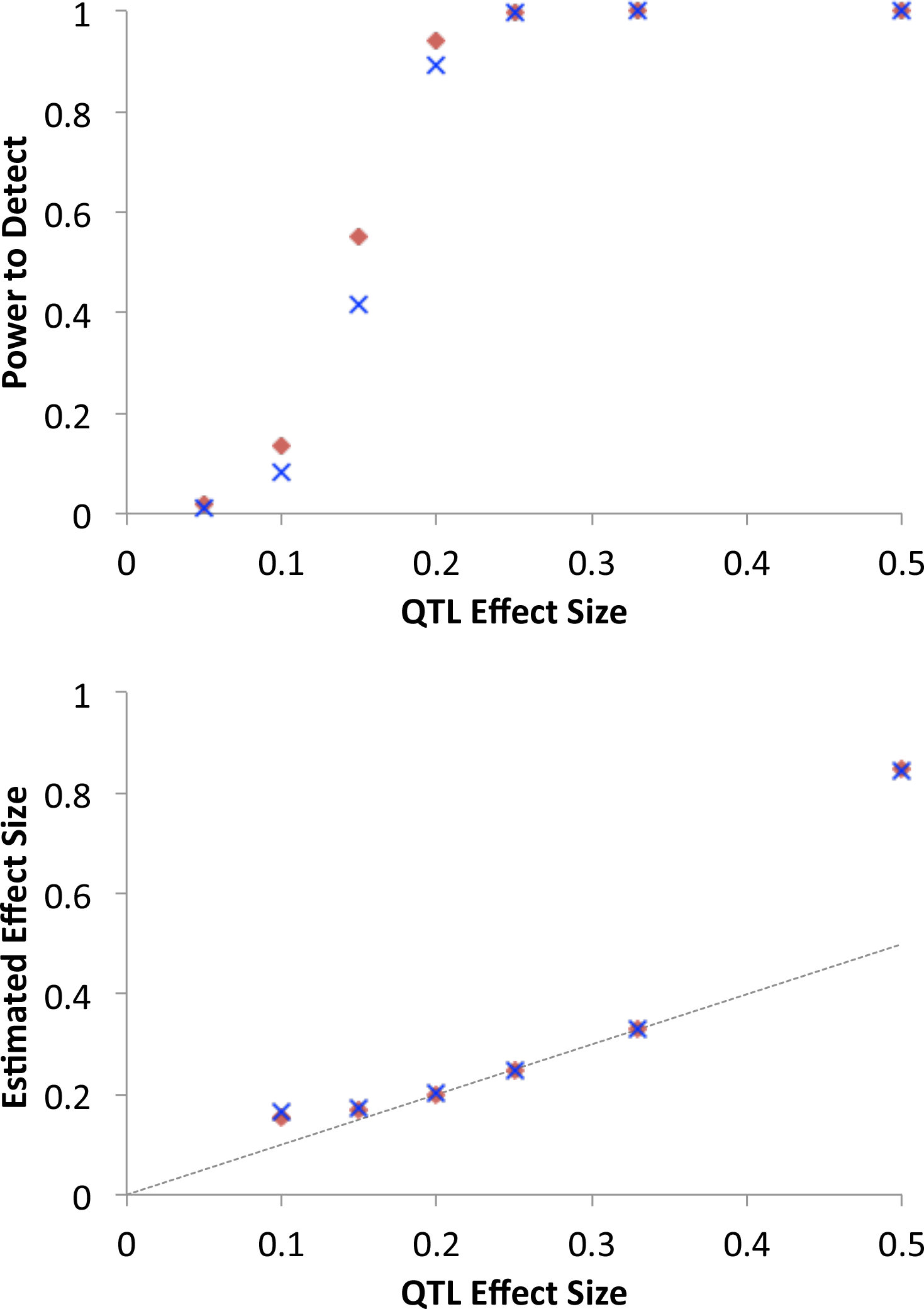
Results of one locus test simulations assessing the performance of the SIBSAM pipline are shown for QTLs on the autosomes (red diamond) and X chromosome (blue X). As shown in the top panel, the scenario investigated here (involving a population of 1,200 individuals with 600 phenotyped after 16 generations and 10% retained in each phenotypic pool) had intermediate power for a QTL explaining 15% of phenotypic variance in the experimental population, with low/high power below/above that mark. As illustrated in the bottom panel and discussed in the text, some upward bias in effect size estimation was observed for the weakest and strongest QTLs examined.

The estimation of QTL strength for significant peaks was quite accurate for intermediate strength QTL (15% to 33%) when the remaining phenotypic variance was random and normally distributed (Figure 7). However, in other scenarios the strength estimate became upwardly biased. For a weaker QTL *(*e.g*.* 10% in this example), there appears to be a “detection bias” in which only the test replicates giving the tallest peaks were deemed significant, and since these peaks are unusually high for *a s* = 10% QTL, their strength was typically overestimated. If strength estimates for non-significant peaks were included, there was no directional bias. The highest QTL strength (50%) also showed upward bias, which may reflect a “saturation effect” of the *a_d_* statistic. Here, peak heights were very close to 1 (individuals were well-sorted into the extreme pools based on QTL genotype), which is the same outcome produced by a QTL with *s* > 50%. Upward strength bias was also observed if the remaining phenotypic variance was produced by other strong QTLs, rather than normally distributed random variance. If a 20% QTL was accompanied by an unlinked 80% QTL (with no environmental/measurement variance), the median estimate of *s* was 24.2%. If a 20% QTL was accompanied by four unlinked QTLs of equal strength, the median estimate of *s* was 31.4% (although power increased from 94% to 100% for both of these cases). In light of the recurrent bias in effect size estimation, the reported quantities are best viewed as rough estimates of QTL strength. Future methodological studies may explore alternative approaches to the estimation of QTL strength in a simulation framework.

Other aspects of SIBSAM inference performed largely as expected on the simulated data. For significant QTL, only around 5% had a false positive secondary peak (in line with null expectations; Figure S2). For QTL strengths with adequate power, approximately the predicted proportion of loci fell within the provided confidence intervals for QTL strength and genomic position (Figure S2), with performance only declining for the weaker *s* = 10% case that was rarely detected for this scenario.

Detection power was also examined for cases involving two linked QTLs (of strength 15% and/or 30%) separated by various distances (2.5 cM, 5 cM, 10 cM, 25 cM). For QTL of equal strength, the 25 cM linkage had no adverse effect on QTL detection. Power was actually slightly higher in the case of 15% QTL separated by 25 cM (relative to the unlinked case), even though 55% of these test replicates had one of the QTLs as a secondary peak. Power to detect a second peak dropped significantly as the distance between QTL dropped to 10 cM and 5 cM (Figure 8). In the case where one QTLs had *s* = 30% and the other had *s* = 15%, power remained high for the stronger QTL at all distances, but was low for weaker QTL at 10 cM or closer (Figure 8).

**Figure 8.**
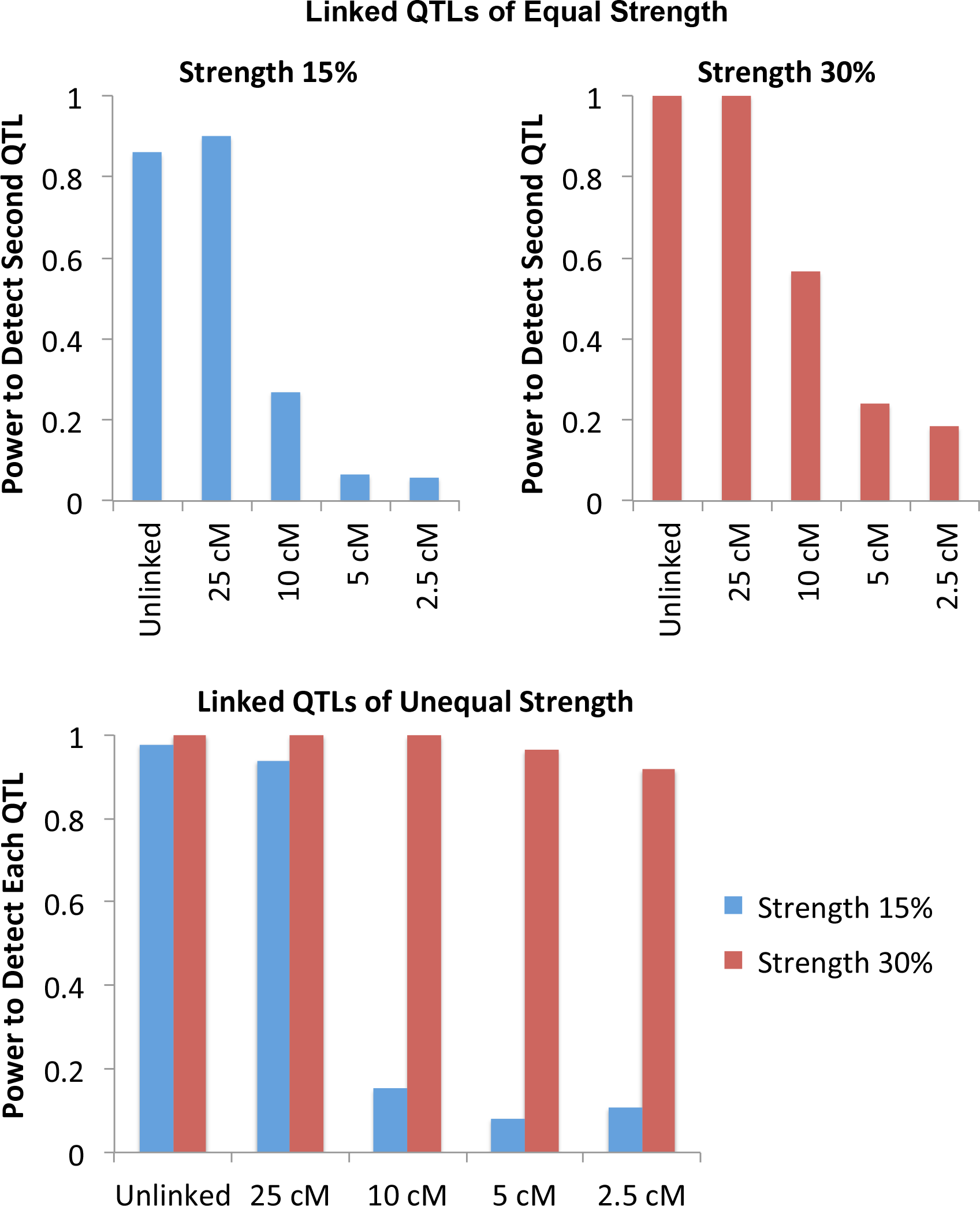
The detection power of SIBSAM in test simulations with two linked QTLs is illustrated. The top panels illustrate the power to detect the second of two linked QTLs of equal magnitude, conditional on detecting the first. The bottom panel illustrates the power to detect either the weaker or the stronger of two linked QTLs of unequal sizes.

## Discussion

Mapping the genetic architecture of phenotypic trait differences remains a challenging but critical problem in the fields of genetics and evolutionary biology. Above, I have compared the behavior of bulk segregant analysis and introgression mapping, while assessing the experimental parameters that modulate their outcomes. I then offered a new simulation-based approach to BSA inference, geared toward systems like *Drosophila* in which hundreds or thousands (but not millions) of individuals can be examined, and in which BSA QTL signals may sometimes overlap each other.

A general principle of QTL mapping is that performance is enhanced by sampling a diverse range of recombinant genotypes. Thus, simulation results suggest that BSA and IM should both be more successful when more generations of interbreeding occur, when larger numbers of individuals are present in the mapping population, and when greater sequencing effort is employed. The importance of sampling at least a few tens of individuals in phenotypically-selected pools is clear as well. These results suggest that the typical method of introgression mapping, in which small numbers of individuals are phenotypically selected every generation or two, is not advisable for mapping oligogenic traits (and is not ideal for monogenic traits either; Figure 4). Instead, if IM is used, larger numbers of phenotyped and retained individuals are desirable. However, based on the criteria employed here, BSA gave a more precise mapping signal than IM for every combination of experimental and QTL parameters examined. This finding may again relate to the principle of maintaining a diversity of recombination breakpoints, which is maximized by avoiding IM's population bottlenecks associated with phenotypic selection during the intermediate generations of interbreeding.

The tradeoffs among BSA, IM, and other mapping approaches are complex and merit further attention. A compelling advantage of BSA is that the same experimental population may be used to map multiple trait differences (*e.g*. once the adults have already reproduced, select for one trait in generation 12, another trait in generation 13, *etc.*). For the same set of experimental parameters as defined here, BSA actually requires less effort than IM during the experiment, since phenotyping must be performed only in the last generation. BSA does require the sequencing of two phenotypic pools (high and low), whereas IM requires just one phenotypic pool to be sequenced (note however that doubling IM depth does not allow it to match BSA's performance; Figure S1). Because both parental strains' genotypes are present across the genomes of mapping population individuals, BSA may be more influenced by the complexities of epistatic interactions.

In the course of a BSA experiment, parental strain ancestry frequencies in the mapping population could deviate from 50%. The effects of genetic drift should be modest when the population size is vastly greater than the number of generations of interbreeding, and SIBSAM allows for drift's occurrence. Although not modeled here, inadvertent laboratory selection could also shift mapping population ancestry frequencies. In general, such ancestry shifts should not lead to false positive QTL, because both phenotypic pools will be equally affected. If ancestry frequencies become extreme, the response of *a_d_* to a QTL could be dampened, leading to reduced power and underestimation of QTL strength. Hence, it may be worthwhile to collect BSA sequence data before an excessive number of generations have elapsed. Genomic regions found to show ancestry shifts could be interesting in their own right, since they may contain drivers of laboratory adaptation, differential mating success, or segregation distortion.

It is more challenging to compare BSA or IM against alternative mapping methods such as those involving individual genotyping (*e.g*. Andolfatto *et al.* 2011) or the generation of recombinant inbred lines (*e.g*. King *et al.* 2012). However, it may be worth evaluating the benefits of combining elements of BSA with these approaches. Following multiple generations in a large mapping population, offspring with extreme phenotypes could be individually genotyped. Or, the mapping population could be used to found a large number of recombinant inbred lines (RILs), with BSA and RIL mapping potentially integrated.

The mapping approach and method described here requires a moderate investment of researcher time and funding, and delivers a range of QTL inferences. While useful in its current form, SIBSAM may also motivate future simulation-based mapping methods. Although motivated by *Drosophila* QTL mapping, this approach may prove broadly useful for non-model insects and other smaller organisms with short generation times.

## Acknowledgements

I thank Karl Broman for helpful discussions regarding SIBSAM and David Begun for suggesting BSA to me several years ago. The UW-Madison Center for High Throughput Computing provided computational assistance and resources for this work. This project was funded by NIH grant R01 GM111797.

## Literature Cited

Andolfatto, P., D., Davison, D., Erezyilmaz, T. T. Hu, J., Mast, et al., 2011 Multiplexed shotgun genotyping for rapid and efficient genetic mapping. Genome Res. 21: 610–617.

Baird, N. A., P. D., Etter, T. S., Atwood, M. C., Currey, A. L., Shiver, et al., 2008 Rapid SNP discovery and genetic mapping using sequenced RAD markers. PLoS ONE 3: e3376.

Bastide, H., J. D., Lange, J. B., Lack, A., Yassin, and J. E., Pool, 2016 Oligogenic adaptation, soft sweeps, and parallel melanic devolution in Drosophila melanogaster. Accompanying manuscript.

Comeron, J., R., Ratnappan, and S., Bailin, 2012 The many landscapes of recombination in Drosophila melanogaster. PLoS Genet. 8: e1002905.

Earley, E. J., and C. D., Jones, 2011 Next-generation mapping of complex traits with phenotype-based selection and introgression. Genetics 189: 1203–1209.

Edwards, M. D., and D. K., Gifford, 2012 High-resolution genetic mapping with pooled sequencing. BMC Bioinformatics 13: S8.

Ehrenreich, I. M., J., Bloom, N. Torabi, X., Wang, Y., Jia, et al., 2012 Genetic architecture of highly complex chemical resistance traits across four yeast strains. PLoS Genet. 8: e1002570.

Haase, N. J., T., Beissinger, C. N., Hirsch, B., Vaillancourt, S., Deshpande, et al., 2015 Shared genomic regions between derivatives of a large segregating population of maize identified using bulked segregant analysis sequencing and traditional linkage analysis. G3 (Bethesda) 5: 1593–1602.

King, E.G., S. J., Macdonald, and A. D., Long, 2012 Properties and power of the Drosophila Synthetic Population Resource for the routine dissection of complex traits. Genetics 191: 935–949.

Lack, J. L., C. M., Cardeno, M. W., Crepeau, W., Taylor, R. B., Corbett-Detig, et al., 2015 The Drosophila genome nexus: a population genomic resource of 623 Drosophila melanogaster genomes, including 197 from a single ancestral range population. Genetics 199: 1229–1241.

Lai, C. Q., J., Leips, W., Zou, J. F., Roberts, K. R., Wollenberg, et al., 2007 Speed-mapping quantitative trait loci using microarrays. Nat. Methods 4: 839–841.

Magwene, P. M., J. H., Willis, and J. K., Kelly, 2011 The statistics of bulk segregant analysis using next generation sequencing. PLoS Comput Biol, 7: e1002255.

Michelmore, R. W., I., Paran, and R. V., Kesseli, 1991 Identification of markers linked to disease-resistance genes by bulked segregant analysis: a rapid method to detect markers in specific genomic regions by using segregating populations. Proc. Natl. Acad. Sci. USA 88: 9828–9832.

Parts, L., F. A., Cubillos, J., Warringer, K., Jain, F., Salinas, et al. 2011 Revealing the genetic structure of a trait by sequencing a population under selection. Genome Res. 21: 1131–1138.

Van Leeuwen, T., P., Demaeght, E. J., Osborne, W., Dermauw, S., Gohlke, et al. 2012 Population bulk segregant mapping uncovers resistance mutations and the mode of action of a chitin synthesis inhibitor in arthropods. Proc. Natl. Acad. Sci. USA 109: 4407–4412.

Wicks, S. R., R. T., Yeh, W. R., Gish, R. H., Waterston, and R. H., Plasterk, 2001 Rapid genemapping in Caenorhabditis elegans using a high density polymorphism map. Nat. Genet. 28: 160–164.

